# Household Transmission Study of Cryptosporidiosis in Bangladesh

**DOI:** 10.1101/269985

**Authors:** Poonum S. Korpe, Carol Gilchrist, Cecelia Burkey, Emtiaz Ahmed, Vikram Madan, Rachel Castillo, Shahnawaz Ahmed, Tuhinur Arju, Masud Alam, Mamun Kabir, William A. Petri, Rashidul Haque, A.S.G. Faruque, Priya Duggal

**Affiliations:** Johns Hopkins University Bloomberg School of Public Health, Baltimore; University of Virginia School of Medicine, Charlottesville; International Centre for Diarrhoeal Disease Research, Dhaka, Bangladesh

## Abstract

**Background:** *Cryptosporidium*, an apicomplexan protozoa, is a leading contributor to diarrheal morbidity and mortality in children under five years old worldwide. As there is no vaccine and no approved drug for *Cryptosporidium spp.* in young children, preventing parasite transmission is crucial. We undertook a pilot case-control study to define the extent of person-to-person transmission of cryptosporidiosis within families in an urban and rural community in Bangladesh.

**Methods:** We enrolled 48 case families with a *Cryptosporidium*-infected child aged 6-18 months. Controls were age-sex matched *Cryptosporidium*-negative children (n=12). Once children were identified, we enrolled all household members. We then followed these individuals for 8 weeks, with weekly surveillance stools and testing with qPCR for *Cryptosporidium spp*.

**Findings:** In the 48 case families, the rate of secondary infections with *Cryptosporidium* was 18.6% (22/118) compared to 0 new infections (0/35) in the 12 control families. In the 22 urban Mirpur households, the secondary attack rate was 30% (18/60) in cases compared to 0% (0/14) in controls (chi-square *p* = 0.018). In contrast, in the 21 rural Mirzapur households, the secondary attack rate was 6.9% (4/58) in case households compared to 0% (0/21) in controls (chi-square *p* = 0.22). Genotyping by gp60 demonstrated infection with the same subspecies in five of six families. Serologic response to *Cryptosporidium* infection was associated with younger age, longer duration of infection, and *C hominis* gp60_IbA9G3R2 infection.

**Interpretation:** The high rate of secondary infection in Mirpur suggests that person-to-person transmission is likely a major source of *Cryptosporidium* infection for young children living in this region. GP 60 genotyping demonstrated direction of infection in 2 households, and concurrent infection in five households. Further work is needed to understand the differences in parasite transmissibility and immunity to different genotypes.

## BACKGROUND

Globally, cryptosporidiosis is a leading cause of diarrhea in young children, and annually it is estimated there are 2.9 to 4.7 million *Cryptosporidium*-attributable cases in children younger than two years of age in sub-Saharan Africa and South Asia (1, 2). Both diarrheal and subclinical *Cryptosporidium* infections have been associated with growth faltering and cognitive deficits (3, 4). Despite the significant public health threat posed by this enteric parasite, there is no effective drug therapy for this patient population and no vaccine.

*Cryptosporidium* is transmitted via fecal-oral spread and outbreaks have been associated with contaminated water supplies (5). In endemic regions, infection is associated with poor sanitation, poverty, contact with domesticated animals, and malnutrition (6). Recent studies aimed at improving household water and sanitation behaviors in Bangladesh showed reduction in overall diarrhea, but they did not result in any decrease in *Cryptosporidium* infections or reduction in stunted growth, a known consequence of *Cryptosporidium* infection (7, 8). In the absence of effective behavioral and environmental interventions and drug therapies, understanding factors involved in transmission and acquisition of infection is imperative.

In prior studies, we have described infection rates as high as 80% with cryptosporidiosis during the first two years of life (9). However, the route of transmission of cryptosporidiosis remains elusive since *Cryptosporidium spp*. can be transmitted through direct or indirect contact (involving contaminated soil, food or water) and subsequent alimentary track infection. Although water contamination is often the cause of *Cryptosporidium* outbreaks we hypothesized that in this setting in Bangladesh, young children were at greatest risk of infection from household contacts, as over 95% of households in this cohort received municipal water, treated by the Dhaka Water and Sewage Authority.

We therefore aimed to understand person-to person transmission in slum communities in Bangladesh by tracking the spread of the disease in the urban and rural households of families with young children. The children in these households were of the age range which is most at risk from *Cryptosporidium* infections. As *Cryptosporidium* infections are common in this population the parasite *gp60* genotype was used to track household transmission.

## METHODS

### Study design

We designed a case-control study of families in urban and rural Bangladesh (Bangladesh Household Transmission Study), nested within a larger birth cohort study on the burden of Cryptosporidiosis in Bangladesh (Cryptosporidiosis and Enteropathogens in Bangladesh; ClinicalTrials.gov: NCT02764918). Prospective longitudinal birth cohorts were established at two sites in Bangladesh. The first in a peri-urban slum community within Mirpur, Dhaka,and the second in a rural subdistrict located 60 km northwest of Dhaka, Mirzapur (Steiner et al, under review). In the parent study, children were enrolled at birth, and followed by twice-weekly home visits for diarrheal illness surveillance.

Children were monitored for the first *Cryptosporidium* infection by monthly surveillance stool collection and testing. Children that reached age six months without infection were continuously screened on a monthly basis for *Cryptosporidium* infection using a rapid immunodiagnostic testin the field (Giardia/Cryptosporidium QUIK CHEK, TECH LAB ®), and results were confirmed using a *Cryptosporidium* specific real time qPCR assay at the icddr,b Parasitology Lab.

Children that tested positive for *Cryptosporidium*, were then consented for enrollment into the Bangladesh Household Transmission Study. All household members, defined as any individual sleeping under the same roof or eating from the same cooking pot, were also consented for enrollment in the study. Written consent was obtained from parents and guardians of all child participants, and directly from all adult participants. Children that tested negative were placed into a pool as potential age- and sex- community matched controls. We enrolled 24 case children and 6 age and sex-matched control children from each site, and their respective household members. We enrolled one matched control for every 4 cases.

### Surveillance

Once enrolled, a baseline demographic and socioeconomic survey was collected from each household. Subsequently, all households were visited weekly for eight weeks, and each enrolled household member completed an illness survey, specifically querying for diarrheal disease in household members. A stool specimen was collected from all household members weekly over eight weeks. A serum sample was obtained from household members at baseline (week 1) and at the end of the study (week 8). GPS coordinates were collected from each household for mapping of infected households.

### Laboratory testing

Stool specimens were collected and transported to the Parasitology Laboratory at icddr.b for testing for *Cryptosporidium* using a multiplex qPCR assay (10). DNA extraction was performed by a modified QiaAmp stool DNA extraction protocol which incorporates a three-minute bead-beating step to lyse *Cryptosporidium* oocysts (Qiagen, Valencia, CA) (10). All specimens testing positive for *Cryptosporidium* were further speciated using the LIB3 assay to distinguish *C. hominis* from non-*C. hominis* isolates (12). The polymorphic region within the gp60 gene was used to genotype *Cryptosporidium* positive samples using previously described methods at the University of Virginia with the modifications of Alves et al (9, 11). Serum samples were tested for the presence of anti-*Cryptosporidium* IgG, using enzyme-linked immunosorbent assay with whole *Cryptosporidium* coated plates (2.5×10*6 oocysts/ml) (WATERBORNE Inc., New Orleans, LA), as described in (13). Anti-human HRP conjugate was used, and absorbance was measured on the ELISA reader at 450 nm.

### Statistical Analysis

A two sample t-test was used to compare differences in means between continuous variables. Chi-square tests were used to compare categorical variables. Multiple linear regression was used to predict association of IgG responses with independent predictors. Statistical analysis was done in Stata 13.1. ArcGIS (10.5.1) software was used for GIS mapping.

## RESULTS

From August 2015 to February 2017, a total of 60 households were enrolled; 24 cases and 6 controls from Mirpur and 24 cases and 6 controls from Mirzapur. This included 60 index children, and 162 household members. Of the household members, 37% were mothers, 16% were fathers, 28% were siblings, 9% were grandmothers, and the remainder (10%) were grandfathers or other relatives.

There were significant differences between households in the two sites (Table 1). The urban site, Mirpur, had a greater percentage of household members testing positive for cryptosporidiosis during the follow up period compared to the rural site, Mirzapur (51% vs 29%, t-test p<0.01). Additionally, there was lower maternal education (mean 4.5 years vs mean 6.4 years p <0.01), more crowding in the home (mean 4 persons/room vs 3 persons/room, p<0.01), and higher occurrence of toilet sharing in Mirpur compared to Mirzapur (mean 3.2 vs 1.1, p = 0.03).

**Table 1.** Characteristics of households from Mirpur and Mirzapur.

|  | Urban Mirpur | Rural Mirzapur | p-value |
| --- | --- | --- | --- |
| Number of case households enrolled | 24 | 24 |  |
| Number of household members enrolled | 75 | 87 |  |
| Number of children under 60 months | 1.2 | 1.2 | 0.5 |
| % of female case children | 61 | 39 | 0.12 |
| % of individuals with *Cryptosporidium* | 51 | 29 | <0.01 |
| Mean years of Maternal Education (SD) | 4.5 (2.7) | 6.4 (4.0) | <0.01 |
| Crowding – Number of persons per room | 4 (0.7) | 3 (1.0) | <0.01 |
| Monthly income (USD) | 229 (208) | 241 (245) | 0.86 |
| % of homes with any animal | 38 | 100 | <0.01 |
| Cow | 0 | 54.2 | <0.01 |
| Goat | 16.7 | 20.8 | <0.01 |
| Chicken/duck | 29.1 | 79.2 | <0.01 |
| Water source | 100% piped | 100% tube well |  |
| % of homes with improved toilet | 79 | 71 | 0.5 |
| Number of households sharing toilet | 3.2 (4.5) | 1.1 (1.0) | 0.03 |

In Mirpur, 18.9% (n= 154) of weekly surveillance stool samples tested positive for *Cryptosporidium.* Of the genotyped samples, 87% were *C. hominis* and 13% were *C. parvum.* In Mirzapur, 10% of weekly surveillance stools tested positive for *Cryptosporidium* (n=94). Of these, 92% were identified as *C. meleagridis*, and 8% identified as *C. hominis*. During the study period, only 14 episodes of diarrhea were recorded, and stool specimens collected (11 in Mirpur and 3 in Mirzapur), and all occurred in index children. Eight of the 14 (57%) diarrheal episodes tested positive for *Cryptosporidium*, 5 in Mirpur and 3 in Mirzapur.

Across both sites there were no infections in the control families. In the 43 case families, the rate of secondary infection was 18.6% (22/118) overall; with a 30% rate (18/60) in Mirpur case households and 6.9% rate in Mirzapur households. In 66% (16/24) of case families in the Mirpur site, multiple family members had concurrent infections with the index child during the follow up period (Figure 2). In 14 of these 24 case families, a second family member who initially tested negative, subsequently became positive during the eight-week follow up period (Figure 2).

Among individuals in Mirpur who tested positive for *Cryptosporidium* during the follow up period, the infants < 2 years had a significantly longer period of shedding (mean 4.1 weeks) compared to individuals over two years (mean 1.7 weeks) (p <0.001). Additionally, there was a higher *Cryptosporidium* parasite burden (lower Ct threshold) in infants compared to those over two years of age (mean Ct thresholds; 26.3 vs 29.0 p= 0.08).

### GP60 genotyping results

In eight of the 24 Mirpur case families, only the index child tested positive for *Cryptosporidium* during the eight-week follow up period, and in six of these families the *Cryptosporidium* isolate was successfully genotyped, the other two had low parasite burden and could not be genotyped. Figure 2 depicts the gp60 genotype of these six children.

In 16 of the 24 Mirpur families, the index child and at least one family member tested positive for *Cryptosporidium* during the follow up period. Four of these families could not be genotyped and in 6 additional families, only the index child was successfully genotyped due to low parasite burden in the other family members. However, in 6 of the 16 families, two or more individuals in the household were successfully genotyped (Figure 2). Figure 2 depicts the genotypes and weeks of infection in these 18 families.

Overall, we detected 12 different *C hominis* genotypes and one *C parvum* genotype. Genotype *C hominis* gp60_1aA18R3 was the most abundant genotype in this cohort (9 samples); This is consistent with a larger study from this region describing nearly 20% prevalence of this genotype (Steiner et al; Gilchrist et al). Additionally, using gp60 genotyping, we detected a temporal pattern to infection with certain genotypes. Genotype *C hominis* gp60_IaA18R3 was present during all three years of the study, whereas gp60_IdA15G1 was only detected in 2015, and gp60_IaA19R3 and gp60_IfA13R3 in 2016 (Table 2).

In families with multiple members genotyped, we observed concurrent infection with the same genotype in four of six families (Families 1, 8, 14,17) and thus we cannot determine directionality of the transmission at entry into the study. In the sixth family (Family 19), the index child is persistently positive for 6 weeks with genotype gp60_IbA9G3R2, while the mother is positive with gp60_IAa18R3 in week one, and never transmits this to the child and does not pick up the child’s genotype despite the persistent shedding. In Family 11, the index child is persistently positive with genotype gp60_IeA11G3T3_V in weeks 2-4 of the study, then in week 5 becomes positive with gp60_1aA19, the same genotype that her five-year old brother is shedding in week 1.

Family 17 is the only family in which *C. parvum* was isolated. In this family, in week 1, both the index child and 6-year-old brother are shedding *C parvum* IIcA5G3R2. The index child has a concurrent infection with *C. hominis* gp60_IbA5G3R3. In week 2, the father becomes infected with *C parvum* IIcA5G3R2, and in week 3, a five-year old brother also becomes infected with the same genotype. The younger brother and father remain infected in week 5, and then in week 6, the mother also becomes infected with the *C parvum* genotype.

### Serologic response

Anti-*cryptosporidium* IgG was measured at week 1 and week 8 in 34 individuals in Mirpur (17 children age two or younger, and 17 subjects older than age two years). There was a greater mean increase in IgG level from week 1 to week 8 in children younger than two years compared to older subjects (mean difference 0.82 vs 0.32, *p*-value = 0.016) (Figure 3). A higher baseline (week1) IgG level was associated with fewer weeks of *Cryptosporidium* positivity during the follow up period when adjusting for age (linear regression; b = −0.85 (95% CI −1.5, −0.18)). The serologic response in week eight was associated with the number of weeks *Cryptosporidium* was detected in stool during the eight-week period when adjusting for age (b= 0.11, 95% CI 0.0, 0.23). Genotype *C hominis* gp60_IbA9G3R2 was significantly associated with a high IgG response when adjusting for weeks of positivity and sex (linear regression; b = 5.37 (95% CI 1.1, 9.6). Genotype *C hominis* gp60_IaA18 was the most common genotype isolated and appeared to be consistently associated with a positive serologic response (Table 2). Genotype gp60_IeA11G3T3 was seen in one index child, and despite the child shedding for four weeks, there was no serologic response to this genotype.

**Table 2.** Subjects with *Cryptosporidium* genotype and *Cryptosporidium* IgG Antibody response. Included in this table are subjects who had stool samples successfully genotyped and had serum IgG results. Children age two years and younger had the most robust increase in IgG from Week 1 to Week 8. In general subjects with greater weeks of stool positivity had the most robust response (index child 3308; index child 3221; index child 3249). *C hominis* gp60_IaA18 was the most commonly detected in this cohort and associated with a positive IgG response. Only one family had infection with *C parvum* gp60_IIcA5G3R2, all individuals tested in this family had a positive serologic response. Index child 3197 tested positive for *C hominis* gp60_Ie_A11G3T3for 4 weeks but failed to show a serologic response.

| Date of enrollment | Family ID | Subject | Age (yrs) | Genotype | Weeks genotype detected | Infection in first 4 weeks | IgG week 8:1 ratio |
| --- | --- | --- | --- | --- | --- | --- | --- |
| Feb 2016 | 19 | index | <1 | *C hominis*<br>gp60_IbA9G3R2 | 6 | 4 | 7.85 |
| Feb 2016 | 13 | index | 1 | *C hominis*<br>gp60_IbA9G3R3_V | 5 | 4 | 4.03 |
| Nov 2015 | 14 | index | <1 | *C hominis*<br>gp60_IaA18 | 5 | 4 | 3.32 |
| Jan 2016 | 1 | index | <1 | *C hominis*<br>gp60_IaA18 | 4 | 4 | 1.29 |
| Jun 2016 | 12 | index | 1 | *C hominis*<br>gp60_Ie_A11G3T3 | 4 | 4 | 0.95 |
| May 2016 | 17 | index | 1 | *C parvum*<br>gp60_IlcA5G3R2 | 4 | 3 | 1.98 |
| Jan 2016 | 5 | index | 1 | *C hominis*<br>gp60_If_A16G1 | 3 | 4 | 1.34 |
| Nov 2015 | 10 | index | <1 | *C hominis*<br>gp60_IdA14 | 3 | 3 | 1.48 |
| Oct 2015 | 16 | index | <1 | *C hominis*<br>gp60_IaA18 | 3 | 4 | 4.89 |
| May 2016 | 17 | father | 32 | *C parvum*<br>gp60_IlcA5G3R2 | 3 | 2 | 1.41 |
| Jan 2016 | 1 | index | <1 | *C hominis*<br>gp60_IaA17 | 2 | 4 | 1.29 |
| Sep 2015 | 3 | index | 1 | *C hominis*<br>gp60_IaA18 | 2 | 4 | 3.47 |
| Sep 2016 | 8 | index | <1 | *C hominis*<br>gp60_Id15G1 | 2 | 2 | 1.22 |
| May 2016 | 12 | index | 1 | *C hominis*<br>gp60_IfA13G1 | 2 | 4 | 2.69 |
| Nov 2015 | 14 | cousin | 2 | *C hominis*<br>gp60_IaA18 | 2 | 2 | -- |
| May 2016 | 17 | brother | 5 | *C parvum*<br>gp60_IlcA5G3R2 | 2 | 1 | -- |
| Jan 2016 | 1 | brother | 8 | *C hominis*<br>gp60_IaA18 | 1 | 1 | 0.94 |
| Dec 2015 | 2 | index | 1 | *C hominis*<br>gp60_Ie_A11G3T3_V | 1 | 2 | 2.24 |
| Sep 2015 | 4 | index | 1 | *C hominis*<br>gp60_IaA17 | 1 |  | 0.79 |
| Sep 2015 | 4 | index | 1 | *C hominis*<br>gp60_IaA18 | 1 | 3 | 0.79 |
| Sep 2015 | 7 | index | <1 | *C hominis*<br>gp60_IdA15G1 | 1 | 3 | 2.17 |
| Sep 2015 | 8 | mother | 28 | *C hominis*<br>gp60_Id15G1 | 1 | 1 | 1.22 |
| Jun 2016 | 11 | brother | 5 | *C hominis*<br>gp60_IaA19 | 1 | 1 | -- |
| May 2016 | 17 | brother | 6 | *C parvum*<br>gp60_IlcA5G3R2 | 1 | 3 | -- |
| May 2016 | 17 | mother | 24 | *C parvum*<br>gp60_IlcA5G3R2 | 1 | 3 | 1.84 |
| Feb 2016 | 18 | mother | 30 | *C hominis*<br>gp60_IaA18 | 1 | 2 | -- |

## DISCUSSION

This is the first study using molecular diagnostics to evaluate and describe person-to-person transmission of cryptosporidiosis to the subspecies level. Our study describes circulating *Cryptosporidium* genotypes, duration of infection, and development of a serologic response in the setting of subclinical infection.

We describe a high rate of *Cryptosporidium* infection among persons of all ages in Mirpur households, similar to that reported in a study from Brazil, where 39% of households were noted to have a secondary infection in a household contact (14). However, when we applied gp60 genotyping, we were unable to establish directionality of transmission in a majority of cases, due to our study design. We recruited families based on a child already having tested positive for *Cryptosporidium* at the start of the study period. Therefore, all new infections could only be studied in relation to the index child. However, genotyping results demonstrate that the index child was not transmitting infection to other family members. In this cohort, genotyping revealed 13 distinct subspecies, and we noted multiple species and subspecies of *Cryptosporidium* circulating within a household; in six households, multiple individuals were infected with the same *Cryptosporidium* subspecies at entry into the study that could reflect person-to-person transmission. We did note two examples of an older household member, a brother and a father, transmitting a novel *Cryptosporidium* infection to the index child.

Though we were unable to establish directionality of transmission in a majority of households, we did note a high rate of new infections among family member in Mirpur, but not in the rural site of Mirazpur. The low rate of secondary infections in the rural site could be explained by differences in predominant species at the different sites. We have previously reported *Cryptosporidium meleagridis* in 90% of samples collected from Mirzapur, whereas *C. hominis* was the most common species in Mirpur (9)(Steiner et al, *submitted*), and in the present study, 92% of speciated samples from Mirzapur were identified as *C. meleagridis*. Avian species have been identified as the natural reservoir for *C meleagridis*, and zoonotic transmission has been described from domesticated chickens to humans (15). In our study, 6 of the 8 families with *C. meleagridis* infection kept chickens at home, suggesting zoonotic transmission. In contrast, the majority of infections in the urban site were attributable to *C. hominis*, for which humans are the only natural host. Thus, the differences in rates of infection may be directly due to different transmission modes of predominant *Cryptosporidium* species in these two different regions. Additionally, it is known that the infectious dose of *Cryptosporidium* can vary from 10 to 1,000 oocysts, depending on the species and strain (16,17). Thus, it is possible that in the Mirpur site, the higher secondary attack rate is directly a result of a lower infectious dose of *C. hominis* compared to the *C. meleagridis* predominant in the rural site.

*C. hominis* gp60_1aA18R3 was the most commonly isolated genotype in this study, and most commonly noted in concurrently infected individuals in the same household. It is difficult to say whether this genotype is more prevalent because it is environmentally ubiquitous, or because it is the most easily transmitted, either due to low infectious dose or other pathogenic characteristics of the parasite.

In two households that were completely genotyped, we observed that infection started in one individual and then spread to other individuals in the household, which we propose is evidence for person-to-person transmission. In one family, we documented transmission from an older sibling to the index child. And in the second family, the only family with *C. parvum* infection, we documented transmission from children to parents and then to other children. Notably, we did not document any other cases of child to adult transmission, presumably because adults had acquired immunity. We have previously documented that *C hominis* is the dominant circulating species in Mirpur, and likely most older subjects have been exposed to *C hominis* at some point in their lives; in contrast *C parvum* has previously been documented in only 2% of infections in this area (Steiner et al, *submitted*). One possibility is that the *C parvum* genotype isolated in the present study was newer to this area, and potentially, even adults had not previously been exposed or developed immunity. This would explain why the *C parvum* IIcA5G3R2 genotype was so easily transmitted between family members of all ages in this household.

Our study suggests that the first *Cryptosporidium* infection may be the most important in stimulating a serologic response, as children with their first infection had the most robust increase in IgG over the eight-week follow up period compared to older subjects. This is consistent with a study from India which showed that IgG response peaked 9 weeks after *Cryptosporidium* diarrheal episode in children. Furthermore, while prior studies have reported an immune response in children after symptomatic infection, our findings are novel as we detected a serologic response even in subclinical, non-diarrheal infections (18,19).

While we did not have records on whether the other household members had ever tested positive for *Cryptosporidium* prior to this study, we do know from our prior work that 77% of children living in this community are infected with *Cryptosporidium* by age two (9); thus presumably, the majority of older children and adults have experienced infection earlier in life. We also noted that higher IgG level at baseline was associated with fewer weeks of *Cryptosporidium* positivity, implying that the elevated IgG may be correlated with an effective immune response. In subjects older than two, the average duration of shedding was significantly lower compared to younger children. This likely reflects greater immunity, and even if older individuals become infected, they may be better able to control infection and have less shedding of the parasite, as reflected by the lower parasite burden we found in older individuals.

Prior studies have noted immunologic cross-reactivity with infection from different *Cryptosporidium* species. One study by Allison et al in Bangladesh demonstrated serologic response to a *C parvum* gp15 antigen in children with symptomatic *C hominis* disease (20). In our study, we used whole *C parvum* oocyst as the antigen in the ELISA, and even though a majority of infections in our cohort were due to *C hominis*, we did note a robust IgG response in children infected with most *C hominis* subtypes. *C. hominis* gp60_IbA9G3R2 was associated with a particularly strong response. Conversely, in a case with *C hominis* gp60_IeA11G3T3, despite persistent infection for 4 weeks, the child failed to demonstrate a serologic response. Our results suggest that there may be species and subspecies level cross-reactivity in immunogenic responses in children which warrants further study with a larger cohort and longer period of observation.

Furthermore, during this period of intense surveillance, we only detected 14 episodes diarrhea, and 8 of 14 were Cryptosporidium-associated diarrhea, and they all occurred in index children. This is consistent with our prior studies which found high rates of non-diarrheal *Cryptosporidium* infection in Bangladesh (9; Korpe et al, *submitted*). We did not find an association of diarrheal infection and transmission, rather in both families with documented transmission, infections were subclinical. This highlights the fact that in households with small children, multiple household members have subclinical *Cryptosporidium* infection, and even though they lack symptoms, they may be silently shedding the parasite and spreading infection.

Limitations of this study include the sample size. This was a pilot study nested within a larger birth cohort. We successfully demonstrated that adult and children family members within this community could be intensively and longitudinally studied, and our findings can now be followed up in a larger study. Secondly, were unable to genotype all the *Cryptosporidium* positive stool samples likely due to the low parasite burden in some stool specimens, primarily in adults. Lastly, we lacked sampling and testing of environmental samples, such as drinking water, soil around latrines, which could have helped to establish environmental burden.

Our study offers unique insight into the endemic transmission of cryptosporidiosis, a disease without vaccine or effective therapy in children. Cryptosporidiosis is known to be spread fecal-orally, and our findings of higher prevalence in areas with more overcrowding and toilet sharing are consistent with this paradigm. The recent WASH Benefits Bangladesh Trial, a cluster-randomized study of clean water, handwashing, and sanitation intervention, did not find any decrease in cryptosporidiosis in children receiving improved water and sanitation (8). Another study from India also failed to show reduction in cryptosporidiosis with clean drinking water (21). Coupled with these results, our findings suggest that even if water and sanitation interventions have high adherence within households, children are still acquiring cryptosporidiosis likely from other infected individuals. In anthroponotic species of *Cryptosporidium*, humans are the only natural host, thus further work is needed to understand determinants of transmissibility of the parasite between persons, including whether age and acquired immunity impact oocyst shedding and infectivity. Ultimately, tackling cryptosporidiosis in endemic regions will require an improved understanding of routes of transmission to young children, as well as concerted efforts at identifying environmental reservoirs of the parasites, and understanding differences in pathogenicity of different species and subspecies.

**Figure 1.**
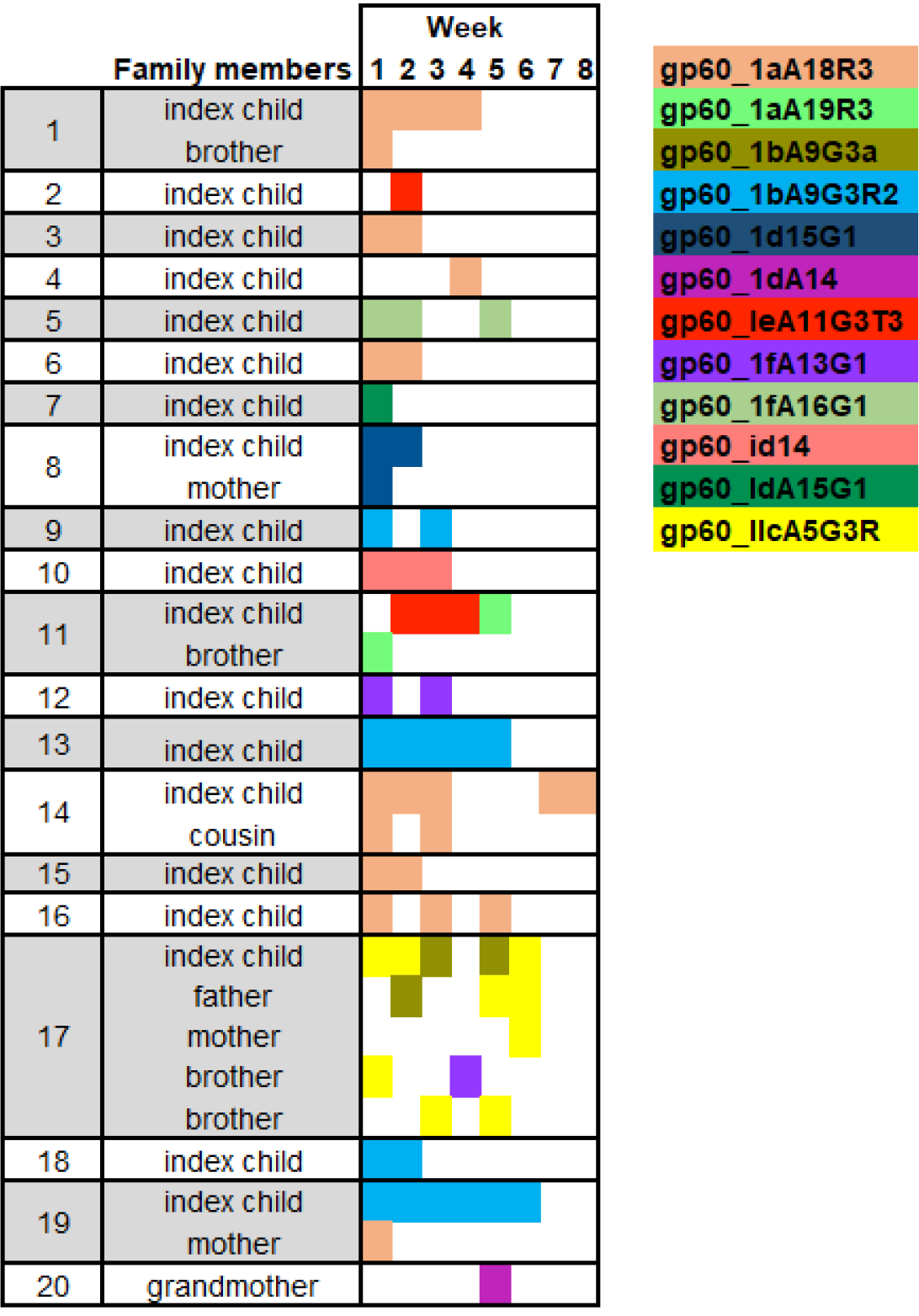
*Cryptosporidium* genotypes detected in 20 Mirpur households over the eight week follow up period. A box shaded in color indicates that individual tested positive for *Cryptosporidiu*m in that week. In 6 of these families, household members were already infected with *Cryptosporidium* at baseline. C hominis gp60_1aA18R3 was the most abundant genotype. Family #11 and #17 demonstrate transmission of a novel genotype from a brother and father to the index child.

